# Unbiased, Cell-free Profiling of Single Influenza Genomes at High-throughput

**DOI:** 10.1101/2024.02.03.578479

**Authors:** Thomas W. Cowell, Wendy Puryear, Chih-Lin Chen, Ruihua Ding, Jonathan Runstadler, Hee-Sun Han

**Affiliations:** Department of Chemistry, University of Illinois at Urbana-Champaign, 505 South Matthews Ave, Urbana, Illinois 61801, USA; Carl R. Woese Institute for Genomic Biology, University of Illinois at Urbana-Champaign, 1206 W Gregory Dr., Urbana, Illinois 61801, USA; Cummings School of Veterinary Medicine, Tufts University, 200 Westboro Rd., North Grafton, Massachusetts 01536, USA

## Abstract

The segmented structure of the Influenza A virus (IAV) genome facilitates reassortment, segment exchange during co-infection. When divergent strains mix across human, agricultural, and wildlife reservoirs novel strains are generated, which has been the source of pandemics. Due to the limited throughput and infection-based assays, IAV reassortment studies has been limited to permissive reassortment. We have developed DE-flowSVP to achieve extremely high throughput, direct profiling of as many as 10^5^ IAV particles in a single-day experiment and enabled quantitative profiling of reassortment propensity between divergent strains for the first time. By profiling reassortants between two naturally circulating low-pathogenicity avian IAVs, we confirmed that molecular incompatibility yields strong preference toward within-strain mixing. Surprisingly, we revealed that two-to-three particle aggregation contributed primarily to genome mixing (75-99%), suggesting that aggregation mediated by sialic acid binding by viral surface proteins provides a secondary pathway to genome mixing while avoiding the co-packaging fitness cost. We showed that genome mixing is sensitively dependent on co-infection timing, relative segment abundances, and viral surface-protein background. DE-flowSVP enables large-scale survey of reassortment potential among the broad diversity of IAV strains informing pandemic strain emergence.

---

Despite its persistent impacts on human health and agriculture through seasonal outbreaks and multiple historical pandemics, Influenza A Virus (IAV) remains challenging to anticipate.^1,2^ IAV undergoes consistent antigenic drift, punctuated by sudden antigenic shifts through reassortment of its segmented genome.^3^ During coinfection, IAV can rapidly exchange its viral RNA (vRNA) segments, producing progeny with mixed genotypes.^4^ Reassortment is the source of novel strains which are difficult to predict.^5,6^ Recent attention has focused on the variety of IAVs that exist across agricultural and wild animal reservoirs, which provide the virus with an array of genetic diversity and potential hosts to reassort and evade existing immunity.^7–10^ Over the course of the past century, reassortment within animal IAVs has caused four major pandemics and many outbreaks in humans, livestock, as well as domestic and wild animals.^11–14^ Currently, there is not a sufficient understanding of genome mixing during reassortment to anticipate potential reassortants before they emerge.^1,2^ To make fundamental progress towards predicting reassortment events, there is a need for large-scale systematic surveys that can assess the reassortment propensity of naturally occurring strains and quantitatively assess the parameters that control genome mixing,^8,9^ but the technology is lacking.

Early studies of reassortment established that incompatibilities exist between differing strains which causes segments to associate more strongly within strains.^15,16^ Because plaque-purifications limits throughput to at most a few hundred individual isolates, studies of rare reassortment events are challenging.^17^ To circumvent this limitation, genetic engineering was applied to remove segment incompatibilities and maximize the reassortment rate. Specifically, silent mutations were introduced to produce two phenotypically identical strains with genotypic markers for the parentage of segments.^18^ This removes the fitness costs associated with segment compatibility allowing for permissive reassortment.^19,20^ While not representative of most natural genome mixing, the important roles of replication incompetent particles to facilitate reassortment through genomic complementation was further identified.^21,22^ This approach was extended to achieve the first assessment of the segment associations during reassortment between two human IAVs (H1N1 and H3N2) that were engineered them to have matched fitness.^23^ However, extending this approach to more divergent strains is not straightforward.

By modifying a single cell RNA-sequencing workflow to target the vRNAs from FACS-sorted infection-positive cells, a >10 fold increase in throughput relative to plaque-isolation methods was achieved.^24^ From sequencing of 37,492 cells, 4,952 complete single virus genomes were detected during reassortment between strains with a history of co-adaptation and successful reassortment.^25^ This approach enabled a thorough investigation of segment associations among the reassortant progeny, quantitatively profiling the composition of the different reassortant genotypes produced upon co-infection.^24^

Notably, the throughput of these approaches is still insufficient to probe reassortment events between more divergent strains. During viral assembly, segment interactions mediate genome assembly.^26^ These packing signals are not conserved across the many naturally circulating IAVs, especially among IAV strains adapted to different hosts, inducing low rates of reassortment.^4^ However, reassortment events between divergent strains are the ultimate source of large antigenic shifts in IAV and pandemic strains.^2,27,28^ For example, the 2009 swine flu pandemic is thought to have occurred through the mixing of avian, swine, and human IAV genes.^12,29^

Also, unbiased profiling is critical to reliable population statistics. However, existing methods rely on cell infection-based assays, where viruses are not measured directly but through infection of a secondary host. Measured abundances will be biased towards genotypes that are best adapted to that host and fail to detect IAV particles that cannot independently self-replicate.^30^ Particularly, defective interfering and semi-infectious particles are replication incompetent, but contribute to reassortment through local complementation.^22,31^ A direct detection method is needed for unbiased profiling of the diversity of particle types present in an IAV population.

Obtaining measurements directly from single virus particles poses a unique analytical challenge, due to the low genomic contents, the requirement of quantitative detection without purification, and the necessity of high throughput. To tackle these challenges, we developed a drop-based microfluidic platform capable of quantitative detection of genome segments from single viruses. Using the new platform, we profiled rare reassortment events between two naturally circulating strains of avian influenza. By quantitatively profiling the genotypes of 10^5^ single virus particles we could detect very low rates of reassortment well below what has been previously possible.^20^ Unexpectedly, we observed 1-10% of virus particle as aggregates that enable mixed genotypes to propagate between incompatible strains by avoiding co-packaging fitness costs. Our analysis of viral progeny revealed that aggregation dominates genome mixing across all infection timings. Further the level of genome mixing sensitively depends on segment abundances, the viral protein background, and the timing of infection events. A database search shows broader distribution of H3 across subtypes when compared to H9, which is consistent with H3’s increased affinity towards sialic acid (SA)^32^ and our observations of increased genome mixing through aggregation for H3N8.

## Results

### Unique Challenges of Cell-free IAV Genome Detection

Widely used assays for single-cell profiling attach barcodes to RNA from cells loaded into droplets.^33–35^ However, in-drop barcoding cannot be directly applied to single IAV particles, due to the low detection efficiency. The IAV genome exists as a single copy meaning a per-molecule detection efficiency of 5-15%^36^ would fail to cover multiple segments. Droplet-based methods for single-molecule detection achieve absolute quantification in drops through enzymatic amplification of a target.^37,38^ However, the required purification step disrupts the particle-level associations needed for single virus measurements. Whole IAV particle loading is likely to lower detection efficiencies as the nucleic acids are not in an accessible form. Natively, each RNA segment interacts with multiple proteins forming a viral ribonucleoprotein (vRNP) complex.^39^

Our approach combines high-resolution and high-sensitivity methods. The first step encapsulates the whole virus into drops, releases the genome from binding proteins, and performs enzymatic amplification in the same compartment. We found that the viral medium showed significant RT-PCR inhibition (Figure S1A) which could be improved through purification using gradient ultra-centrifugation (UC). However, even UC-purified viruses still showed reduced segment counts relative to a purified control (Figure 1A) measured using a ddPCR method (Figure S1B-C). We hypothesized that inhibition was caused by binding proteins that prevent reverse transcription and designed a protease-lysis condition compatible with downstream enzymatic amplification to test this. The lysis buffer consists of a PCR-compatible detergent and a thermolabile protease. After genome release, the protease is heat inactivated and RT-PCR reagents are added. This method enabled near-quantitative detection of the viral RNA (Figure 1A). We also confirmed that viruses are intact by treating each sample with RNAse.

**Figure 1:**
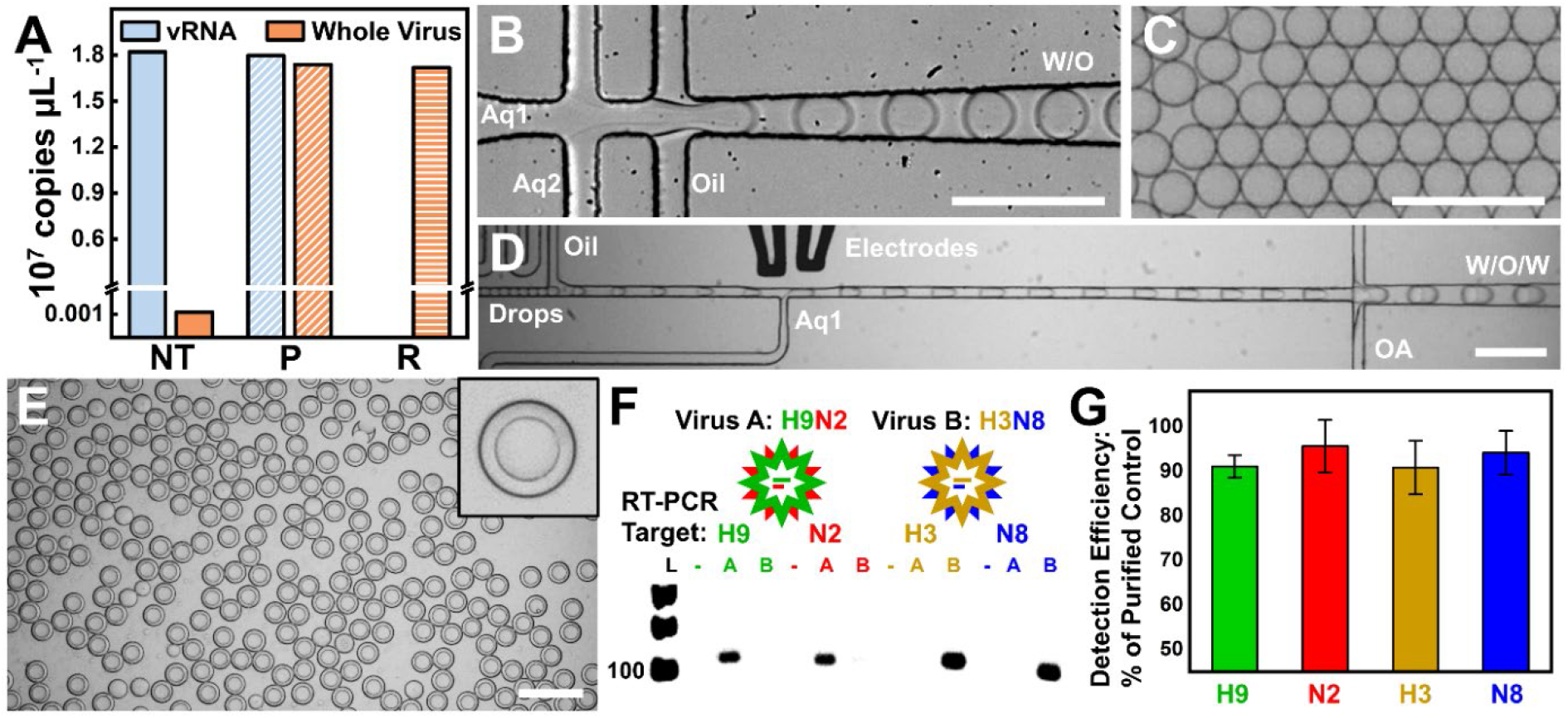
A) Summary of ddPCR experiments used to develop the drop compatible IAV detection assay. Segment copy number is compared between purified viral RNA (vRNA) and intact particles (Whole Virus) by ddPCR after NT: No treatment (solid fill), P: protease treatment (diagonal lines), or R: RNAse treatment then protease treatment (horizontal lines). B) Drop generation device during co-encapsulation of single IAV particles and proteolysis buffer. C) Monodisperse water-in-oil (W/O) drops containing single virus genomes after heat inactivation of the protease. Scale bars (B-C) are 100 μm. D) Integrated picoinjector and DE dropmaker device during operation. Close-packed drops from C are respaced in oil for reagent addition by Aq1. The picoinjected drops are then further dispersed in the OA phase and collected as a water-in-oil-in-water (W/O/W) double emulsion. E) Monodisperse DE drops produced after picoinjection. Scale bars (D-E) are 200 μm. F) Four color TaqMan assay targeting HA-NA segments from selected IAV strains. Gel electrophoresis confirms the correct amplicon is generated for each assay. G) Quantification of RNA counts obtained by droplet processing of single viruses compared against purified RNA controls. Detection of each segment is achieved with upwards of 90% efficiency.

The challenge of multiplexing the PCR step was addressed using TaqMan probes. We could readily achieve four-color detection which can identify the parental origin of two segments. Given that the viral surface proteins, hemagglutinin (HA) and neuraminidase (NA) determine the viral subtype, we chose to target these segments in naturally circulating IAVs isolated from the environmental reservoir. During PCR, each amplicon produces a unique fluorescent color encoding the genotype of the virus. We leveraged the field of double emulsion (DE) microfluidics to achieve drop analysis using flow cytometry (∼10^6^/hour).^40–42^ Per our previous study, DEs allow confident detection of rare events due to their coalescence properties. We combine in-drop IAV lysis with multi-segment RT-PCR and DE flow cytometry to realize direct single IAV profiling at high throughput.

### DE-flowSVP Workflow

The overarching strategy of DE-flowSVP is illustrated in Scheme 1. Direct counting of original strains, incomplete viruses, and reassortants provides an unbiased and quantitative measure of viral populations at single virus resolution. To validate the platform and highlight its utility for systematic reassortment profiling, we investigated genome mixing between two strains of low-pathogenicity avian IAVs; H9N2 (A/northern shoveler/Interior Alaska/10BM16764R0/ 2010) and H3N8 (A/mallard/Interior Alaska/10BM11415R0/2010). These naturally circulating viruses were isolated from different waterfowl species within the same region and timeframe. Reassortment between these subtypes could yield H3N2 and/or H9N8. Based on previous observations and isolated reported into the Bacterial and Viral Bioinformatics Resource Center (BV-BRC), H3N2 is reported frequently while H9N8 is rarely observed, but there is no history of reassortment between these strains.

DE-flowSVP integrates two microfluidic devices and flow cytometry analysis. The first device is a dropmaker, that rapidly compartmentalizes single virus particles into droplets while introducing the lysis buffer (Figure 1B). These single virus drops (Figure 1C) are incubated for genome release and protease inactivation. At this stage, the droplets now contain accessible RNA genomes. We developed a modified picoinjector device^43^ to form a DE at the outlet (Figure S2A) by implementing new flow-confinement strategies^44^ (Figure S2B-C). Figure 1D shows this device in operation as it adds a controlled volume of RT-PCR reagents to each and forms a DE from the result. Since we collect as a DE (Figure 1E), PCR-induced coalescence does not result in merging of the inner compartments, fully maintaining the single virus resolution during signal generation.^45^ Drops remain monodisperse throughout the workflow (Figure S3).

Using a multiplex TaqMan assay (Figure 1F), the presence of H9, H3, N2, or N8 segments in each drop is encoded as a four-color fluorescent signal. These on and off fluorescence signals in each drop can be readily distinguished during imaging or flow cytometry (Figure S4). After gating for correctly sized DE drops, the four-parameter data is projected onto two-parameter slices and the (2^4^) color combinations are measured at high throughput (∼10^6^ drops/hour). Across three replicates, the per-segment detection efficiency of DE-flowSVP was measured as >90% by comparing purified RNA to whole viruses (Figure 1G).

### Validation of Single Virus Resolution and Quantification of Segment Dropouts

To validate the single virus particle resolution, a mixed sample of H9N2 and H3N8 virus isolates (1:1) was measured by DE-flowSVP at a loading of 1 in 10 drops (Figure 2A). Flow detection reveals the genome contents of each drop. Most events were negative for all segments (N) indicating empty drops. The remaining positive drops are counted and plotted on a pie chart (Figure 2B). Using a model of random doublets, these raw drop counts are corrected to account for chance co-encapsulation events. After doublet correction, the genotypes of 25,542 single virus particles were measured. The expected virus strains were most abundant and very few mixed strain particles were detected confirming single virus resolution.

**Figure 2.**
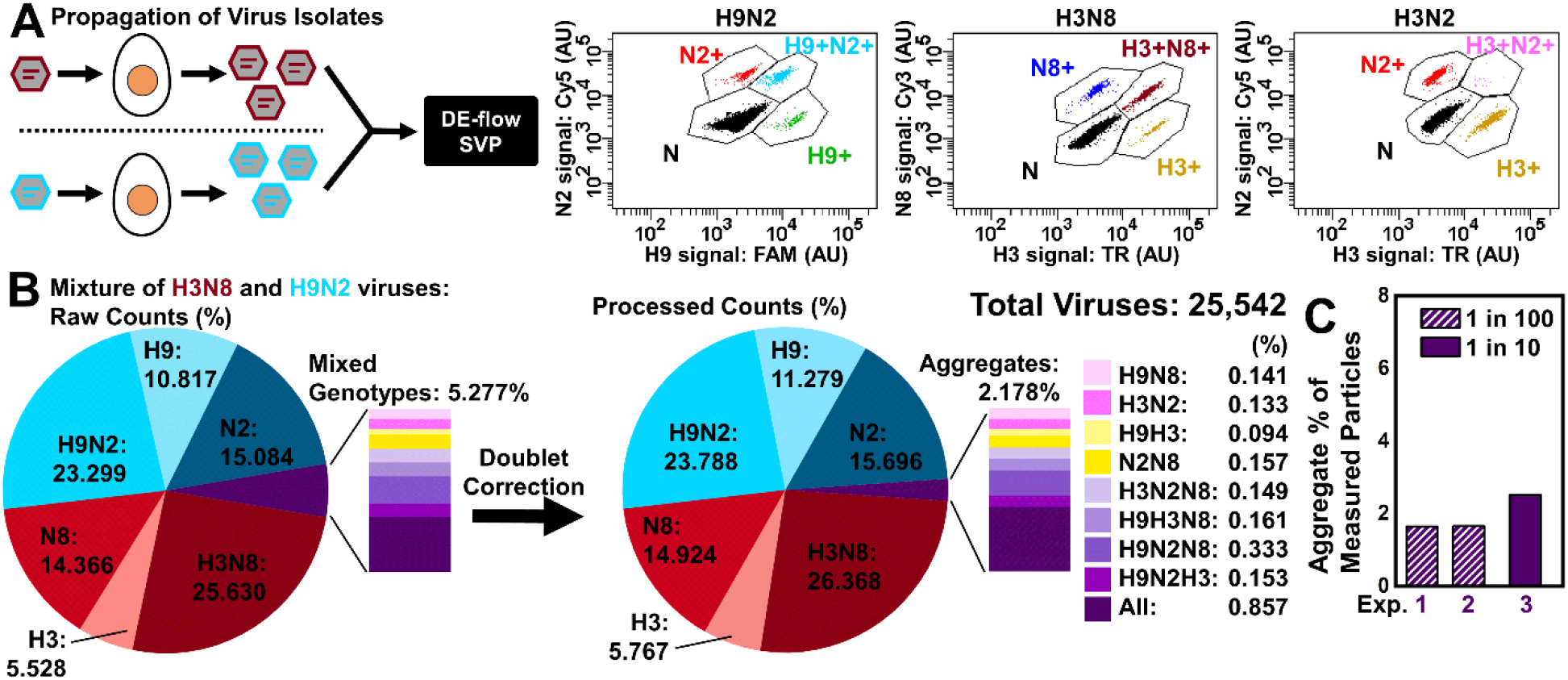
A) Diagram of strain mixing experiment to validate single virus resolution. Pure H3N8 and H9N2 viruses are cultivated in isolation, then mixed and processed by the DE droplet flow cytometry workflow. Raw FACS counts show droplet populations primarily correspond to the original pure virus strains. B) Summary of raw single virus counts, and processed counts that correct for double encapsulation events. Notably some particle aggregates are detected. C) Aggregate measurements from three technical replicates at various drop loadings. If random co-loading of particles is the source of mixed genotypes, a 10-fold dilution of their concentration should result in a ∼100-fold reduction in co-detection which was not the case.

DE-flowSVP also enabled the first high-throughput profiling of defective or incomplete IAV particles. The measured rates of segment dropouts, which can be due to packaging failure or deletions spanning the TaqMan amplicon, are presented in Figure S5A. Viruses featuring segment dropouts are challenging to study as they are unable to replicate. While they have been measured using FISH-imaging to co-localize segments, this approach is low throughput.^46^ The measured dropout rates varied from 11.3% - 21.7%, which is consistent with previous studies if not slightly higher.^46,47^ This small difference in measured dropout rates could be explained by a strain difference or as an effect of the host.

### Quantitative Profiling of Single Virus Aggregates

Interestingly, a small fraction of the measured particles from the species mixing experiment had genome segments from both strains (Figure 2B). Because each virus was propagated in isolation, we would not expect these segment combinations to exist. Three technical replicates were performed at different particle loadings (1-10%) and ∼2% of particles with mixed genotypes were consistently observed (Figure 2C). The consistent frequency of mixed genotypes indicates that is not measurement error but suggestive of IAV particle aggregates. It is documented that IAV populations can self-aggregate through HA binding of sialic acid residues displayed on the viral membrane.^48^ The activity of NA to cleave these groups assists in dissociation of viral aggregates.^48,49^ To test if the observed mixed genotypes result from aggregation, we modeled them as aggregates with each two-particle combination occurring at the conditional frequency of the single particle components (Figure S5B). Because ddPCR cannot distinguish 1 template copy from 2 or more copies of the same template, only the mixed genotype aggregates are considered. The correlation is reasonably good (R = 0.83) confirming that these observations are consistent with viral aggregation. It further indicates that aggregation occurs independent of the genome sequences packaged within particles. Our measurement is consistent with previous studies using FISH or TEM imaging that have revealed IAVs aggregate at a rate of 1-20% with >90% of these aggregates containing 2 or 3 single viruses.^46,50^ Aggregates have been shown to be biologically functional by increasing infection multiplicity which can impact infection dynamics including reassortment.^30,31^ Our new platform facilitates systematic profiling of viral aggregates, which opens doors to study how these populations impact infection dynamics and evolution.

### Systematic Profiling of Genome Mixing between Natural Stains H9N2 and H3N8

To profile the reassortment potential of two natural strains, embryonated chicken eggs (ECEs) were selected as a model host, permissively infected by avian influenza and representative of vertebrate infection with an immune system.^51,52^ To simulate natural co-infection events we performed simultaneous or sequential exposure (Figure 3A-B). The viral progeny was purified by ultracentrifugation and measured with DE-flowSVP. In total, drop processing takes around 5 hours, and flow analysis an additional 2-3 hours allowing for ∼10^5^ single viruses to be profiled in a single-day experiment.

**Figure 3.**
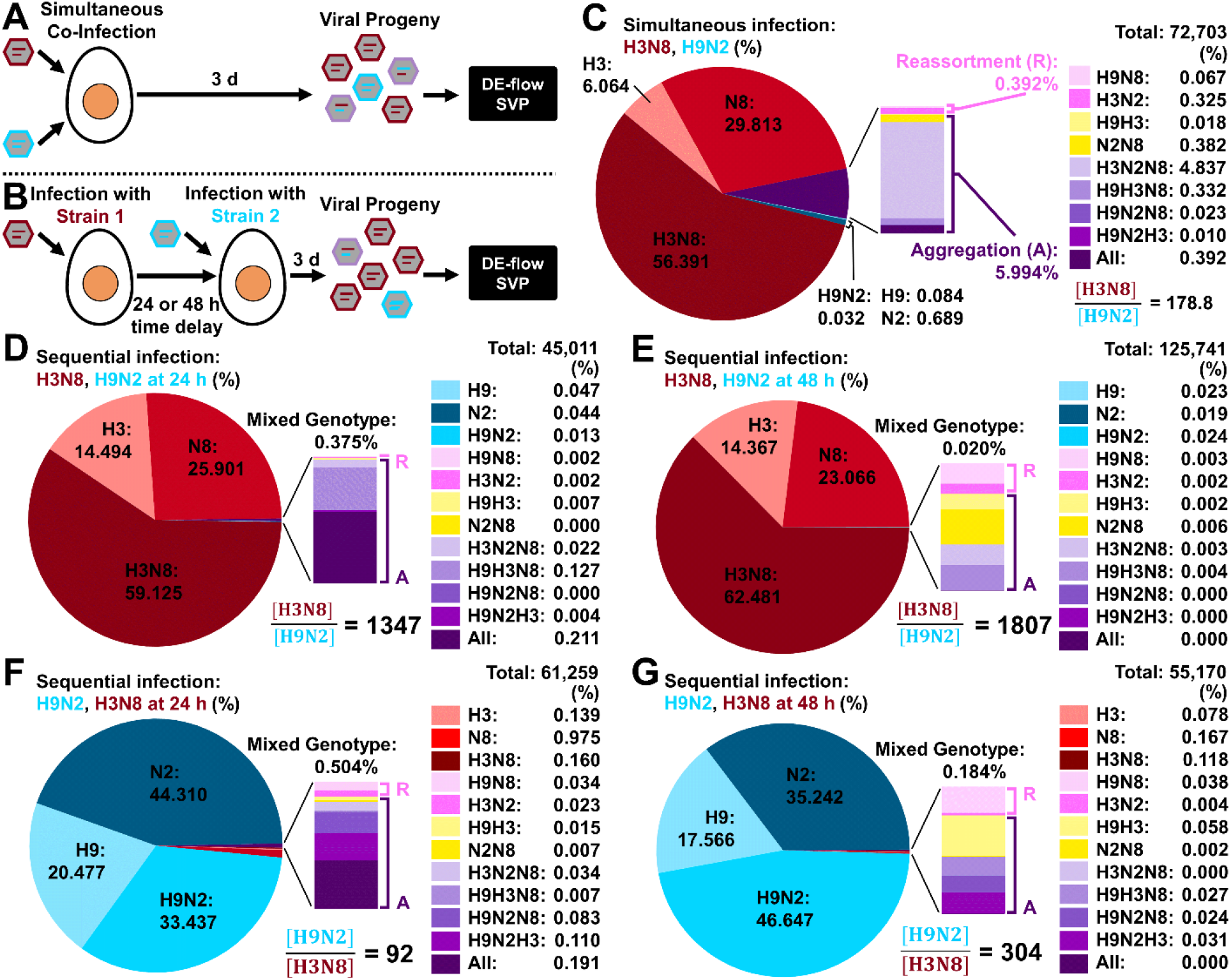
Design of experiment to systematically profile genome mixing between H3N8 and H9N2. Viruses are either simultaneously co-infected (A) or sequentially infected (B) with a time delay of 24 or 48 h. Progeny are collected 3-days post infection and UC-purified for detection by DE-flowSVP. C) Summary of progeny virions produced by simultaneous co-infection. Summary of viral progeny measurements when H3N8 is infected first followed by H9N2 at 24 (D) or 48 (E) hours later. Summary of the progeny viruses produced when H9N2 is allowed to infect first followed by H3N8 at 24 (F) or 48 (G) hours.

This throughput enhancement enabled us to perform the first systematic profiling of reassortment between two unrelated strains from the natural reservoir. When co-infected simultaneously, the segment abundances in the progeny indicate the relative fitness of each virus within the culture system. As expected from divergent strains, there was a large difference in relative fitness that yielded a 178.8-fold bias towards H3N8 segments in the viral progeny (Figure 3C). Since DE-ddPCR eliminates merging, we can confidently detect rare events as low as 0.01% of the population corresponding to seven droplets. From simultaneous co-infection, 6.386% of the progeny displayed a mixed genotype. While H9N8 and H3N2 are valid reassorted virus-types, these particles could also form through aggregation. We define aggregates as the particles with two copies of at least one targeted segment from different parental strains, which assigns the maximum contribution to reassortment. Accordingly, 94% of genome mixing in this sample was facilitated through viral aggregation.

We next delayed the inoculation of the second strain by 24 or 48 hours and measured the progeny as before. The genotypic profile produced when H3N8 is inoculated first is summarized in Figure 3D-E. Notably, we observe a reduction in genome mixing to 0.375 and 0.02% with a 99% and 75% contribution from aggregation as well as significant bias in segment abundance (a factor of ∼1350 and ∼1800) at 24 and 28 hours respectively. When the strain of lower fitness, H9N2, was administered first (Figure 3F-G) the bias in segment abundance skewed by ∼92 and ∼300 towards H9N2 and the rate of mixed genotype progeny fell to 0.504% and 0.184% (87% and 77% contribution from aggregates) at 24 and 48 hours respectively. Across all samples the ratio of HA to NA segments within strains was close to 1 except for simultaneous co-infection where genome mixing was highest (Table S1). In that sample, N2 was almost 7-fold higher among the progeny relative to H9. Having higher abundance for N2 compared to H9 in the progeny could be explained by a higher fitness of H3N2 compared to H9N8, which matches with their abundances in the BV-BRC database, although those reported strains are distinct from those studied here.

### Parameters Governing Genome Mixing

From this systematic profiling, multiple determinants of genome mixing can be identified. Across all samples, the formation of mixed genotype particles between these viruses was driven primarily through aggregation (Figure 4A). This result is reasonable as co-packaged reassortants have an additional fitness cost that limits their formation as we observe in Figure 4B. When accounting for segment abundance, the association within strain was always greater indicating segment incompatibilities that place a strong fitness cost on viruses that package segments across strains.

**Figure 4.**
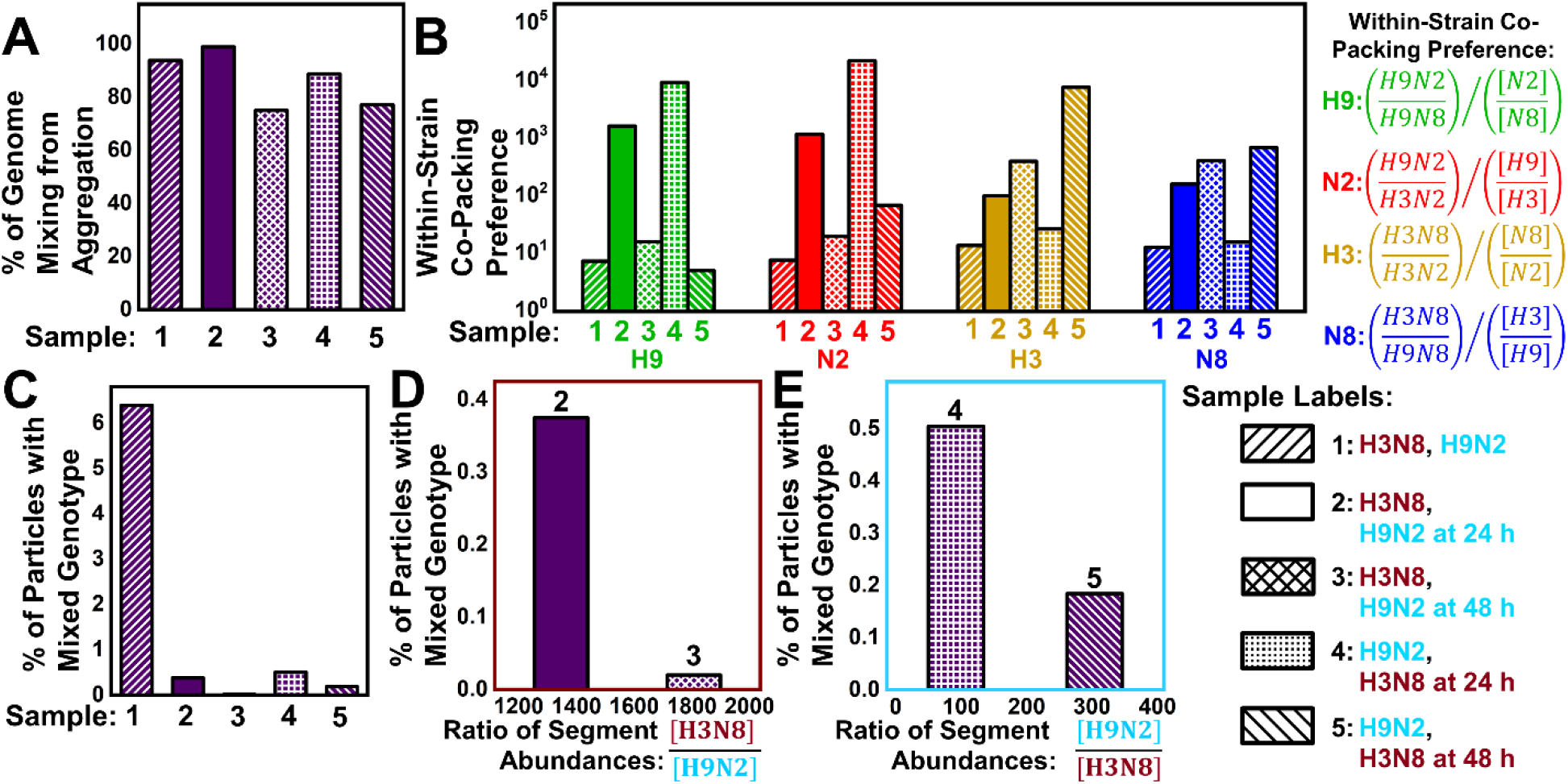
Sample Labels: 1: H3N8, H9N2, 2: H3N8, H9N2 at 24 h, 3: H3N8, H9N2 at 48 h, 4: H9N2, H3N8 at 24 h, 5: H9N2, H3N8 at 48 h. A) The % of genome mixing from aggregation for each sample. Aggregation contributes a substantial majority of all genome mixing. B) Direct experimental evidence of the RNA-RNA co-packaging network. Across all segments and sample types, co-packaging within-strains is preferred. C) The % of viral particles with a mixed genotype across sample conditions. Simultaneous co-infection shows significantly higher genome mixing. During co-infection where H3N8 segments (D) or H9N2 segments (E) are dominant, the likelihood of genome mixing decreases with increasing bias in segment abundance.

Another parameter that influences genome mixing is the co-infection timing. Longer delays between infection events will result in an increasing skew in segment abundance across strains and a lower level of genome mixing. This result is straightforward as only aggregation or co-packaging of segments from different strains can yield mixed genotypes. Our observations of a decrease in genome mixing with the infection interval (Figure 4D-E) aligns with previous measurements of reassortment using permissive strains with matched fitness.^18^ In this previous study, the limited throughput (<100 progeny viruses) prevented detection of reassortants after delays of 24 hours or more even in the absence of segment incompatibility.^18^ Due to the enhanced throughput, we can extend the time-scale of reassortment studies to include multi-day delays which better reflect the natural course of infection and confirm that genome mixing can occur even with large time gaps at reduced probability.

It is notable that genome mixing occurred at least ten-fold more often during simultaneous co-infection than any sequential infection event (Figure 4C). Although the 24 hour delayed H3N8 sample showed a similar level of segment bias as simultaneous co-infection (∼179 vs ∼92), simultaneous infection showed ∼12-fold higher frequency of mixed genotype progeny. This result suggests that the exposure of different strains to each other during their initial stages of replication promotes greater mixing. This likely occurs through increased self-aggregation mediated by a higher density of uncleaved SA-groups present on the surface of freshly produced viruses.^53^

Protein background also has a substantial effect on genome mixing (Figure 4D-E). Since the segment abundances correlate with viral protein expression, the protein background is expected to be primarily H3N8 for samples 2-3 (Figure 4D) and H9N2 for samples 4-5 (Figure 4E). Straightforwardly, genome mixing decreases with increasing segment bias when the protein background is held constant. Notably, the rate of genome mixing was similar for the 24 hour delayed samples 4 (∼0.4 and ∼0.5%) despite significantly different fold-biases (∼1350 vs ∼92). Since genome mixing was driven mainly through surface-protein induced aggregation, this suggests that the H9N2 protein background is less conducive to aggregation.

## Discussion

Here, we developed a platform to release single IAV genomes in drops for multiplexed quantitative TaqMan assay. DE-flow cytometry enables multi-parameter drop screening at high-throughput to measure as many as 10^5^ individual particles in a single experiment and quantitatively profile reassortment events. By combining multiple experiments, the throughput could be pushed to achieve >10^6^ virus particles. DE-flowSVP is readily adaptable to other IAV strains by designing PCR primers. Since HA and NA segments feature high sequence diversity, unique TaqMan assays are easily achievable for reassortment studies of almost all strain pairs. It may also be possible to distinguish the parental origin of segments with low sequence diversity as TaqMan probes with LNA-bases have been used to detect SNPs.^54,55^ Improved multiplexing of the platform may be achievable through additional TaqMan probes which could cover more segments during single virus profiling. However, further multiplexing is potentially challenging as dye and quencher pairs are limited for Taqman assays and PCR competition may impede signal generation. Although we focus on IAV here, the new platform can be applied to study reassortment more broadly across segmented viruses that infect all branches of the tree of life or even to probe multiple loci within a single genome segment.^56^

DE-flowSVP is unique among approaches to study IAV as it directly counts viral particles which allows for definitive assessment of environmental, molecular, or culture parameters. Previous methods have relied on cell-infection to profile single viruses, which inevitably biases the measured population towards those variants best adapted to replicate in this host. Since our detection method is cell-free, it is now possible to quantitatively assess culture parameters and investigate their impact on viral progeny. A detailed understanding of culture effects must precede any large-scale systematic reassortment profiling of the circulating strains in environmental reservoirs. DE-flowSVP can be combined with existing RNA-seq technologies to compare the relationship between vRNA expression in cells and distribution of progeny particle genotypes produced by co-infection events. This unique analysis can reveal whether a low segment abundance among viral progeny results from inefficient vRNA production or selective exclusion during co-packaging.

Due to its enhanced throughput, our technique is capable of fundamentally different studies of IAV reassortment. Prior studies have been limited in the number of single viruses measured which has necessitated a focus on reassortment between compatible strains. In permissive reassortment, segment interactions during packaging are most important in determining which viral progeny will propagate.^19,23,24^ However, reassortment between divergent strains is the source of large antigenic shifts that lead to pandemic emergence.^2,20,28^ The extremely high throughput of our technology can address the low frequency of rare reassortants and overcome the highly skewed distribution of progeny resulting from strains with mis-matches fitness. For divergent strains, where significant segment incompatibilities exist, the formation of co-packaged reassortants is limited. Instead, our observations suggest that aggregation can provide a pathway that circumvents the strict molecular interactions that shape co-packaging and allow mixed genotypes to propagate (Scheme 2).

Aggregation is random with respect to the gene contents and is mediated through the viral surface proteins and their specific interactions with SA residues. Aggregates have gained attention recently, due to their role in increasing the infectivity of segmented and multipartite viruses.^57^ Our observations are consistent with prior studies that have found that most IAV self-aggregation results in two or three particle aggregates^50^ which is unlikely to substantially limit transmission through aerosolization.^58^ Although it is known that most IAVs fail to successfully replicate, aggregation increases the probability of success through gene dosing.^21,22,31,58^ Towards genome mixing, aggregates act a means for simultaneous co-infection and allow for a greater diversity of segments to reach the new host while circumventing the strict fitness cost of co-packaging. In a secondary host, such as in an avian-to-mammalian spillover event, unique genetic combinations could display elevated fitness that drives further co-adaptation of incompatible segments.

To investigate this further, we considered which parameters can influence delivery of mixed genotypes. Since aggregation, which is independent of the genome contents of individual particles, drove genome mixing, the relative abundance of the various gene segments determines the likelihood of mixing. Our analysis suggests that when two strains have a larger difference in relative fitness, the sensitivity towards infection timing will be greater due to the effect of segment bias. Additionally, because aggregation is mediated by SA binding, the HA-NA activity of the co-infecting strains is expected to impact genome mixing. The functional balance of HA activity to bind SA residues and NA activity to cleave SA is crucial to effective IAV attachment and release during replication.^59,60^ For example, when the HA gene is replaced to impart increased activity viral self-aggregation increases.^61,62^ Recall that similar levels of genome mixing occurred in the 24h delay samples where a skew in segment abundances were vastly different: ∼1350-fold towards H3N8 and ∼92-fold towards H9N2. This result suggests that the H3N8 protein background produces mixed genotype progeny through aggregation more efficiently in comparison to H9N2. Additionally, prior studies have identified that H3 has higher affinity towards the avian type α2,3-linked SA than that of H9.^32^ This heightened HA activity may explain the increased aggregation for the H3N8 protein background.

If H3 activity induces higher aggregation and an increase in genome mixing, we would expect it to be more broadly distributed across subtypes. Since there is not a standard test to determine the functional balance of HA-NA,^60^ we performed a database search to compare the distribution of H3 vs. H9 and N2 vs. N8 across subtypes within avian hosts. The total number of unique H3 or H9 genome sequences for each subtype reported in the BV-BRC database are presented in Table S2. Since H3N8 and H9N2 are dominant segment pairings among avian strains, these strains are unlikely to result from reassortment and so were excluded from the analysis. There was a clear overabundance of H3 across NA subtypes in both absolute and relative terms, which is consistent with its greater propensity for genome mixing. Performing a similar analysis on N2 and N8 did not reveal a clear pattern across subtypes (Table S3), which may indicate that the observed increase in genome mixing is driven more strongly by HA activity in these cases. It is important to note that these database results are influenced by a variety of factors including natural abundances, host range overlaps, sampling biases, and more. By comparing across all available avian subtypes, we seek to address some of these effects.

The database analysis provides support for surface protein-mediated aggregation to promote genome mixing across subtypes in nature. If confirmed, reassortment events between divergent strains might be anticipated by measuring strain properties such as HA-NA activity and relative fitness.^63,64^ For example, two divergent viral strains with small differences in relative fitness and high HA relative to NA activity would be expected to produce particles with mixed genotypes at higher frequency. This combined parameter of HA-NA activity and relative fitness has the potential to improve risk assessment of strains for their pandemic potential.

As shown, DE-flowSVP can generate new hypotheses and then confirm these predictions through direct quantification of the viral progeny. Moreover, it is now possible to perform systematic screening of the large variety of circulating strains to assess their potential to reassort and identify additional parameters impacting genome mixing. This capability is important as climate change affects the habitats and migration of wildlife impacting the spatial and temporal overlap of reservoir hosts in ways that provide new opportunities for reassortment. The recent global dissemination of high-pathogenicity H5Nx avian influenza viruses has posed significant concerns due to its intense agricultural impact as well as the increased risk of inter-host transmission.^65^ Reassortment of these strains with endemic low-pathogenicity avian influenza viruses provides pathway to introduce greater genetic diversity across hosts.^66^ With this new approach, predicting the emergence of pandemic strains may become feasible through a thorough understanding of which reassortment events are most likely to occur.

## Acknowledgements

We would like to thank the Roy J. Carver Biotechnology Center for their support. T.W.C, C.-L.C., and H.-S.H thank the University of Illinois at Urbana-Champaign for funding through the UIUC Startup Grant and also the National Institute of Health (R01 AI148385) for support. T.W.C. and H.-S.H. further recognize financial support from the Gordon and Betty Moore Foundation through Grant GBMF9195 to the Carl R. Woese Institute of Genomic Biology. W.P. and J.R. acknowledge funding of this work in part through contracts to the Center for Research on Influenza Pathogenesis and Transmission from the National Institute of Allergy and Infectious Diseases CEIRS and CEIRR (contract numbers: HHSN272201400008C, and 75N93021C00014).

## Author Contributions

T.W.C., W.P., J.R., and H.-S.H. designed experiments. T.W.C. and R. D. designed microfluidic devices. T.W.C. and C.-L.C. fabricated devices and performed droplet experiments. W.P. performed virology experiments. T.W.C., W.P., J.R., and H.-S.H prepared the manuscript.

---

**Scheme 1:**
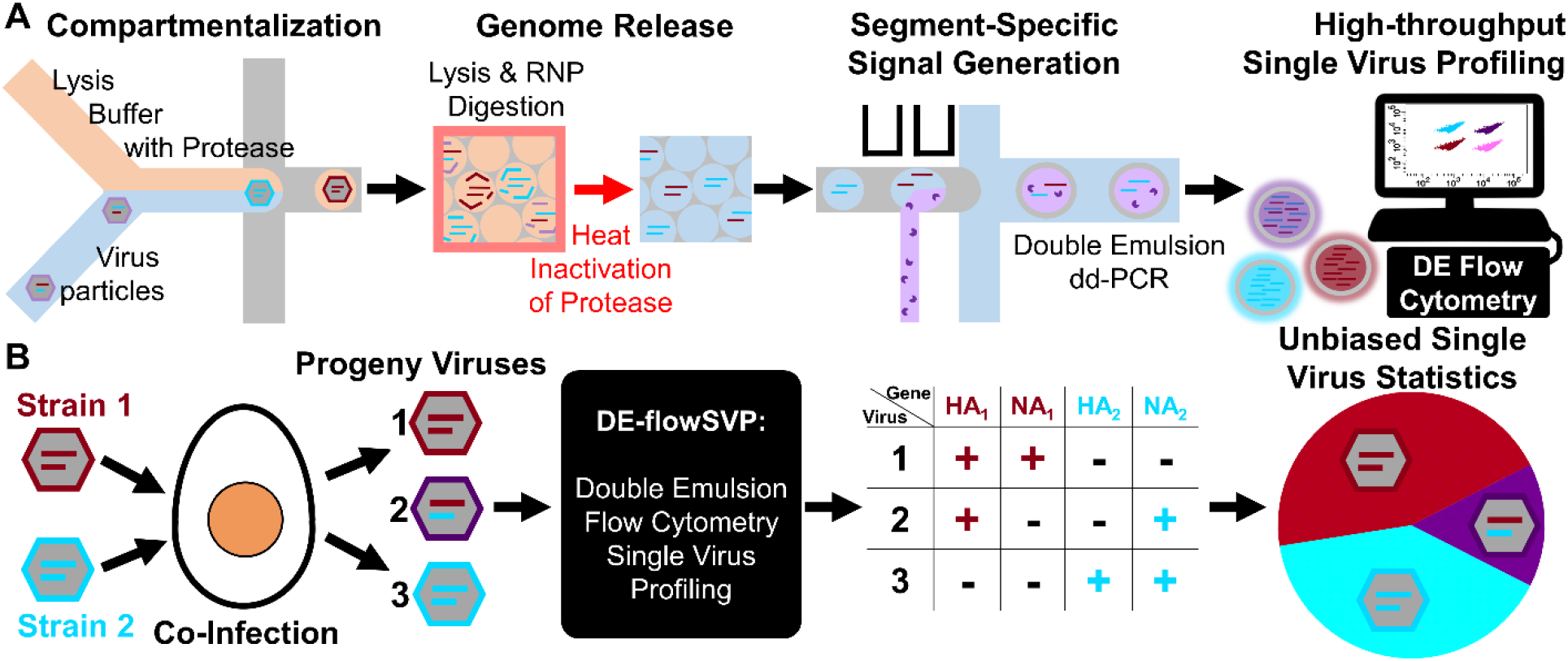
A) Strategy for high-throughput direct genomic profiling of single IAV particles. Individual viruses are compartmentalized into drops with a proteolysis buffer. The viral capsid is lysed and vRNPs are digested by the protease resulting in complete genome release. Thermal inactivation of the protease enables subsequent enzymatic reactions in the same drop. Picoinjection adds RT-PCR TaqMan assay reagents and drops are collected as a double emulsion to enable high-throughput drop detection by flow cytometry. The multi-color readout of each droplet represents the single virus genotype. B) Diagram of reassortment profiling experiment. Two distinct strains co-infect the host producing many progeny virions including reassortants. Using DE flow-SVP, the progeny viruses are directly profiled, and the HA-NA genotype is measured to give the unbiased population statistics produced by the co-infection event.

**Scheme 2.**
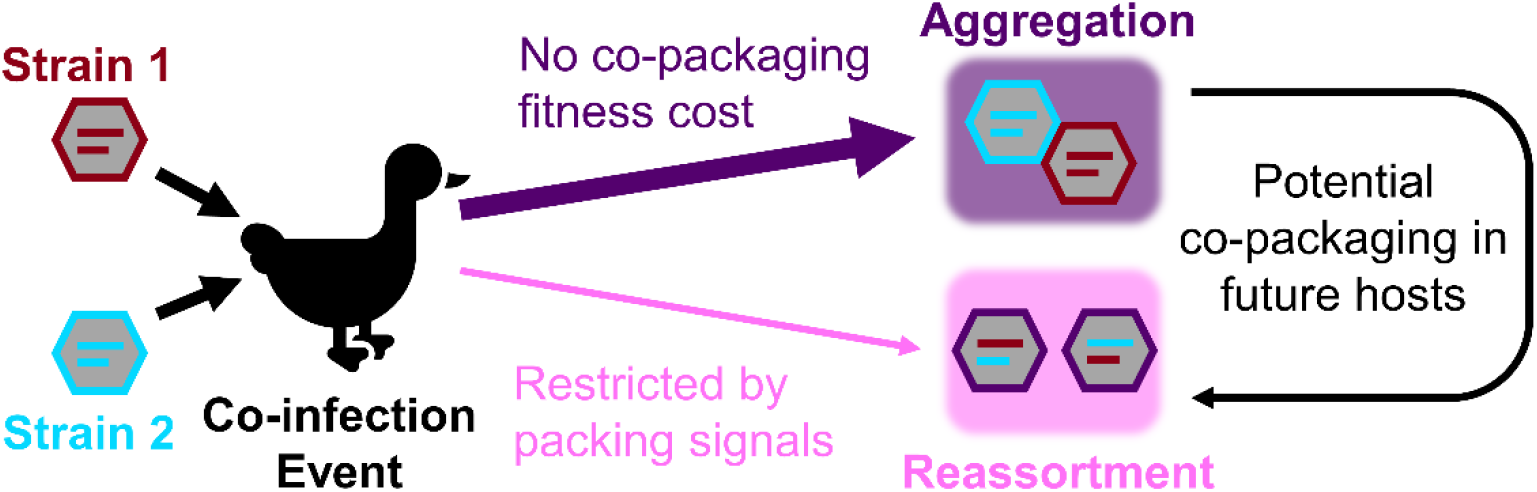
Schematic representation of genome mixing resulting from a co-infection event between two strains of avian influenza. Co-packaging fitness costs restrict reassortment between incompatible strains. Aggregation provides an alternate pathway for the propagation of the mixed genotype. Further selection in subsequent hosts can facilitate co-packaging of reassortants from aggregates.

