## Supplementary material for "Unbiased, Cell-free Profiling of Single Influenza Genomes at High-throughput": SI

**Supporting Information:**

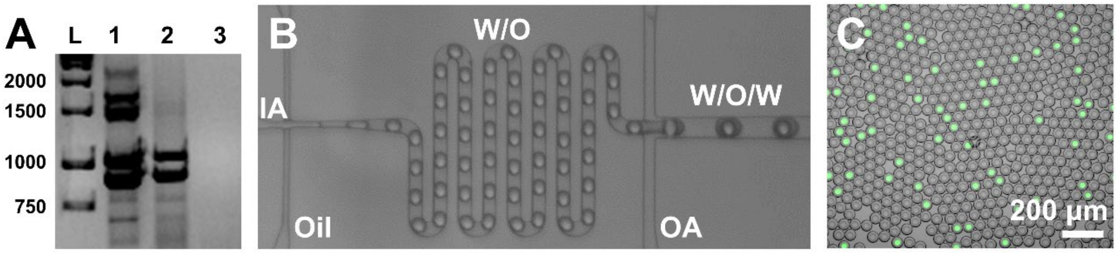


**Figure S1**. A) Gel image showing multi-segment RT-PCR using universal primers. Purified vRNAs amplify efficiently and display the full genome banding pattern (1). When whole viruses collected in amnio-allantoic fluid are added along with purified vRNAs. Inhibition of RT-PCR results in missing segments and lower yield (2). Whole viruses in amnio-allantoic fluid fail to detect due to inhibition of RT-PCR (3). B) Operation of DE dropmaking device during ddPCR quantification of segment copy number. C) Composite image of post-stained amplicons generated during drop digital RT-PCR in double emulsion drops.

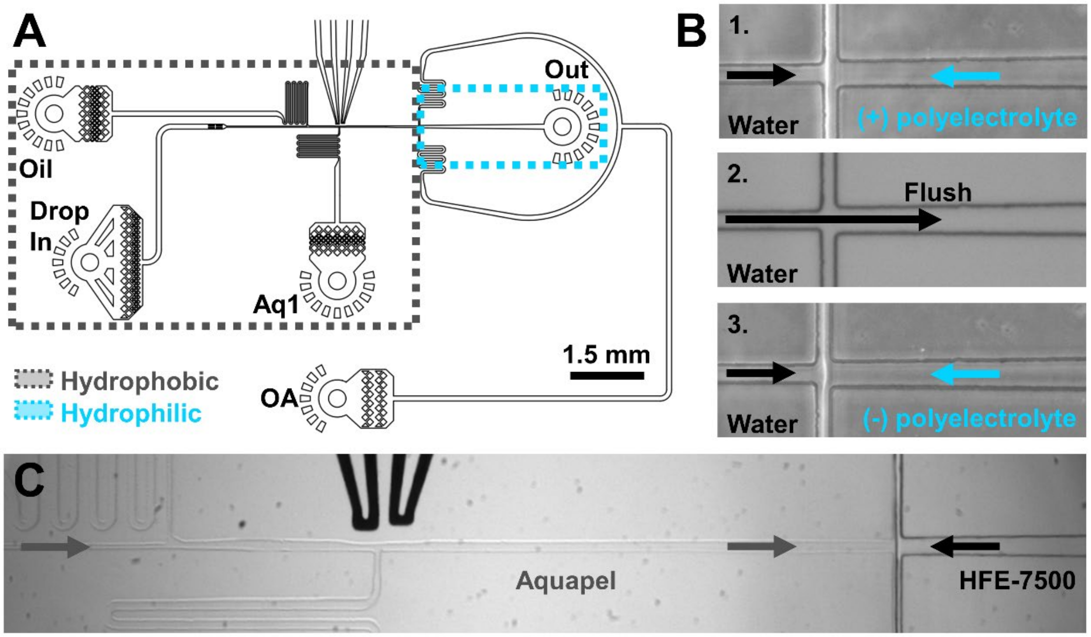


**Figure S2.** A) Device schematic of the integrated picoinjector and DE dropmaker. B) Flow confinement approach to spatial patterning of a poly-electrolyte coating at the DE junction. C) Flow confinement strategy to apply a hydrophobic coating to the single emulsion portion of the device.

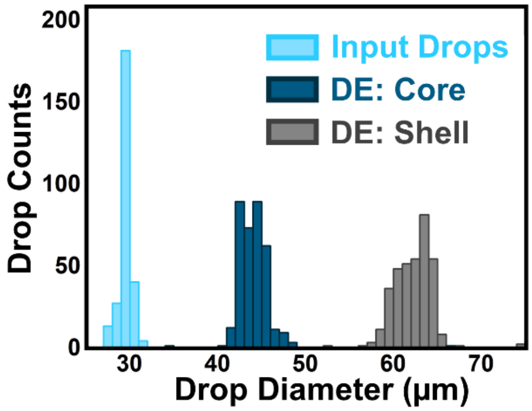


**Figure S3.** Histogram of drop sizes throughout the DE-flowSVP workflow. High quality monodisperse emulsions are generated at each stage. The sizes of more than 200 drops were measured for each sample.

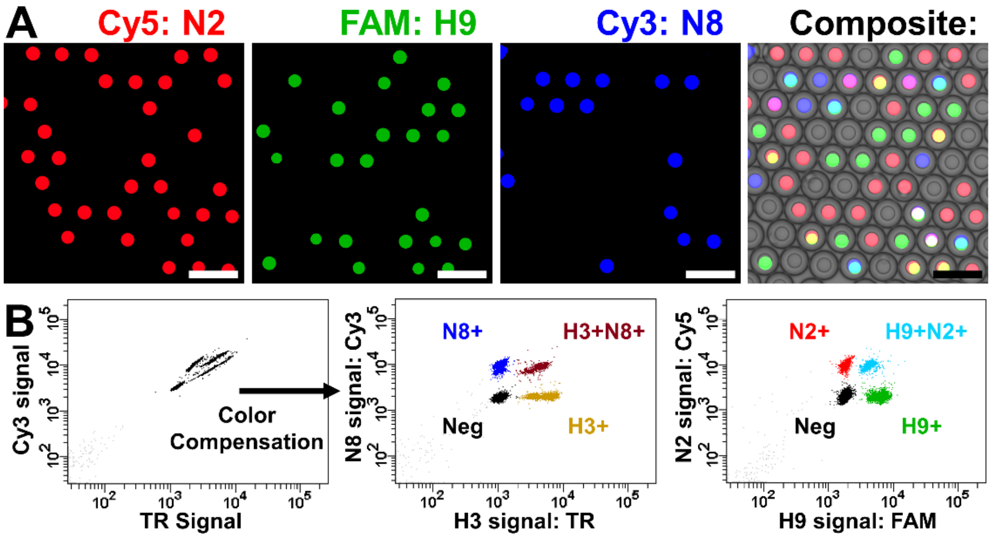


**Figure S4.** A) Microscope imaging of 3-Color DE drops. B) 4-Color DE drop detection using a BD FACSymphonyA1. Following color compensation each positive and negative population for the identification of H3N8, H9N2 are clearly resolved.

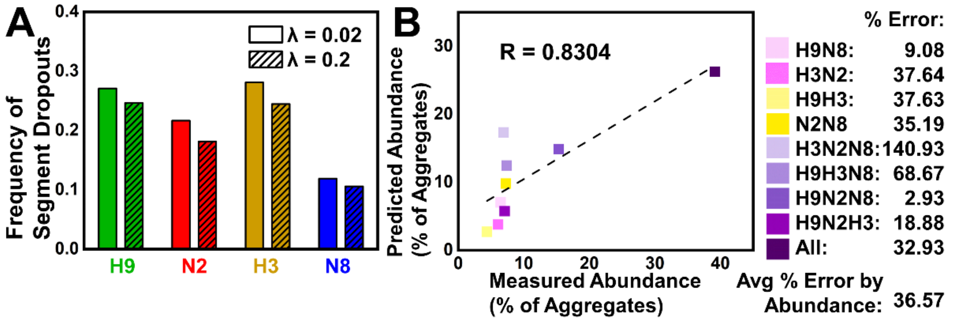


**Figure S5.** A) Measured rates of segment dropouts calculated from species mixing experiments. These particles either fail to package the HA or NA segment for that strain or have large deletions that span the TaqMan assay region resulting in a failure to detect that segment. At higher loadings, there is a slight decrease in the measured dropout rate due to increased aggregation. B) Scatter plot comparison of measured and predicted abundances of each aggregate type. Aggregates were modeled as consisting of two particles from the following particle types; H9, N2, H9N2, H3, N8, H3N8 at their measured abundances.

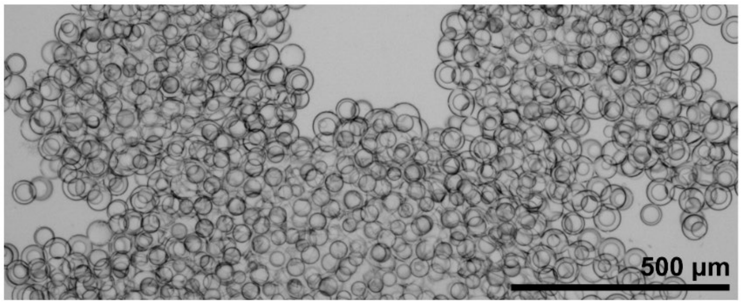


**Figure S6.** Microscope image of DE drops after the occurrence of “BSA Clumping”. Inclusion of ~1-2% tween-20 along with a reduction in denaturation temperatures to 86 °C and post-PCR storage at room temp (>20 °C) is sufficient to prevent drops containing 2% BSA from clumping.

**Table S1.** Relative Segment Abundance within Strains

| Segment Ratio | 1: H3N8, H9N2 | 2: H3N8, H9N2 at 24 h | 3: H3N8, H9N2 at 48 h | 4: H9N2, H3N8 at 24 h | 3: H9N2, H3N8 at 48 h |
|---|---|---|---|---|---|
| N2:H9 | **6.990** | 0.736 | 1.439 | 0.945 | 1.275 |
| N8:H3 | 1.346 | 1.154 | 2.147 | 1.113 | 0.969 |

**Table S2.** Genome sequence abundance in BV-BRC database by NA subtype:

| Avian Hosts | N1 | **N2** | N3 | N4 | N5 | N6 | N7 | **N8** | N9 | Avg. |
|---|---|---|---|---|---|---|---|---|---|---|
| **H3** | **840** | 4,440 | **308** | **54** | 207 | **2,657** | **197** | 16,898 | 120 | 626.1 |
| H9 | 320 | 46,505 | 58 | 23 | **278** | 100 | 85 | 60 | **366** | 175.7 |
| $H3 - H9$ | 520 | NA | 250 | 31 | -71 | 2,557 | 112 | NA | -246 | **450.4** |
| $\frac{(H3 - H9)}{(H3 + H9)}$ (%) | 44.8 | NA | 68.3 | 40.3 | -14.6 | 92.7 | 39.7 | NA | -50.6 | **31.5** |

**Table S3**. Genome sequence abundance in BV-BRC database by HA subtype:

| Avian Hosts | H1 | H2 | **H3** | H4 | H5 | H6 | H7 | H8 | **H9** | H10 | H11 | H12 | H13 | H14 | H15 | H16 | Avg. |
|---|---|---|---|---|---|---|---|---|---|---|---|---|---|---|---|---|---|
| **N8** | 379 | 176 | 16,898 | **2,529** | 7,475 | 2,641 | 340 | 28 | 60 | **1,003** | 317 | 126 | **1,139** | 15 | **8** | **8** | 1156.0 |
| **N2** | **1007** | **829** | 4,440 | 1,680 | **12,600** | **6,933** | **2,467** | 29 | 46,505 | 577 | **1,437** | **177** | 930 | 16 | 3 | 0 | 2048.9 |
| $N8 - N2$ | -628 | -653 | N/A | 849 | -5125 | -4292 | -2127 | -1 | N/A | 426 | -1120 | -51 | 209 | -1 | 5 | 8 | **-892.9** |
| $\frac{(N8 - N2)}{(N8 + N2)}$ (%) | -45.3 | -65.0 | N/A | 20.2 | -25.5 | -44.8 | -75.8 | -1.8 | N/A | 27.0 | -63.9 | -16.8 | 10.1 | -3.2 | 45.5 | 100.0 | **-10.0** |

**Supplementary Video 1.** Operation of the integrated picoinjector and DE dropmaker as it controllably adds reagents to droplets containing accessible single virus genomes.
